# Dichloroacetate reverses sepsis-induced hepatic metabolic dysfunction

**DOI:** 10.1101/2020.06.09.142430

**Authors:** Rabina Mainali, Manal Zabalawi, David Long, Nancy Buechler, Ellen Quillen, Chia-Chi Key, Xuewei Zhu, John S. Parks, Cristina Furdui, Peter W. Stacpoole, Jennifer Martinez, Charles E. McCall, Matthew A Quinn

## Abstract

Dramatic metabolic reprogramming between an anabolic resistance and catabolic tolerance state occurs within the immune system in response to systemic infection with the sepsis syndrome. While metabolic tissues such as the liver are subject to end-organ damage during sepsis and are the primary cause of sepsis death, how their metabolic and energy reprogramming during sepsis state ensures survival is unclear. Employing comprehensive metabolomic screening, targeted lipidomic screening, and transcriptional profiling in a mouse model of septic shock, we show that hepatocyte lipid metabolism, mitochondrial TCA energetics, and redox balance are significantly reprogramed after cecal ligation and puncture (CLP). We identify increases in TCA cycle metabolites citrate, cis-aconitate, and itaconate with reduced fumarate and triglyceride accumulation in septic hepatocytes. Transcription analysis of liver tissue supports and extends the hepatocyte findings. Strikingly, the administration of the pyruvate dehydrogenase kinase (PDK) inhibitor dichloroacetate (DCA) reverses dysregulated hepatocyte metabolism and mitochondrial dysfunction. Our data indicate sepsis promotes hepatic metabolic dysfunction. Furthermore, our data indicate that targeting the mitochondrial PDC/PDK energy homeostat rebalances transcriptional and metabolic manifestations of sepsis within the liver.

---

Sepsis is a potentially life-threatening condition that occurs due to the body’s overwhelming response to an infection. This reaction leads to aberrant immune responses and if not diagnosed and treated early after its onset, may limit survival by inducing coagulopathy, altered microvasculature, and dysregulation of the host’s metabolism and organ function^1–7^. Sepsis accounts for one in three hospital deaths in the U.S. and millions of deaths each year globally, highlighting its hazard to public health^8,9^. The high mortality rate associated with sepsis reflects the lack of a clinically viable molecular-based therapeutic target. Therefore, understanding the pathogenesis of sepsis at both the molecular and organismal level is of utmost importance to address major gaps in knowledge regarding sepsis.

The response to sepsis has classically been characterized as a biphasic phenomenon, where the acute phase is characterized by high energy consumption and hyperinflammation associated with oxidative stress, followed by cellular reprogramming to a low energy, anti-inflammatory state of immunometabolic paralysis, with accompanying organ failure^8,10–12^. Additionally, during the early phase of sepsis increased catabolism of fats, proteins, and carbohydrates, associated with high rates of oxygen consumption and ATP synthesis is observed^13–16^. Following the acute phase is a hypometabolic state where ATP production and mitochondrial respiration decreases^13^. It is postulated that this is a protective mechanism to overall lower the metabolic demands of the cell and help with its recovery^1,17,18^.

Of particular interest to us, during the hyper-inflammatory anabolic phase of sepsis, an increase in the expression and activity of pyruvate dehydrogenase kinase 1 (PDK1) consistently occurs^19,20^. This enzyme is one of four PDK isoforms that reversibly phosphorylates serine residues on pyruvate dehydrogenase complex (PDC) E1a subunit, inhibiting the conversion of pyruvate to acetyl coenzyme A (acetyl CoA)^21^. Inhibition of this important enzymatic activity that connects glycolysis to tricarboxylic acid (TCA) cycle, oxidative phosphorylation (OXPHOS), and the lipogenic pathway is thought to be an important mechanism driving the dysfunction of mitochondrial respiration and cell bioenergetics observed during sepsis^19,22^. Therefore, PDK serves as a potential therapeutic target as it can allow for the downstream oxidation of glucose to continue, restoring OXPHOS and mitochondrial functioning that we know to be altered in both immune and non-immune cells during sepsis^16,19,23,24^.

The liver is an important metabolic and immune organ due to its role in nutrient metabolism and production of acute phase proteins. However, our understanding of transcriptional alterations and subsequent metabolic manifestations elicited by sepsis still remain limited^16,25–28^. Furthermore, the effects of DCA at restoring hepatic metabolic function in the context of sepsis is completely unknown. Hence, we set out to characterize the hepatic manifestations of sepsis, with the overall goal of identifying global metabolic pathways subject to dysregulation and whether these pathways are restored by DCA treatment.

## Results

### Sepsis Impairs Hepatic Mitochondrial Metabolism

To test the long-term hepatic transcriptional changes elicited by sepsis, we performed RNA-seq in whole livers thirty-hours post cecal ligation puncture (CLP). At this time point, septic mice exhibit tolerance, in which immunometaboic paralysis and end organ dysfunctions decrease survival. Ingenuity pathway analysis (IPA) of the top physiological pathways subject to alteration in response to sepsis revealed a significant increase in the acute phase response pathway, highlighting a persistent inflammatory state in the liver into the chronic phase of sepsis (Fig. 1a). Of particular interest were the findings that oxidative phosphorylation and mitochondrial dysfunction were the top two enriched pathways in the liver of septic mice (Fig. 1a). To gain further insight into the effects of sepsis on the transcriptional regulation of mitochondrial function, we performed gene set enrichment analysis (GSEA) of the oxidative phosphorylation pathway (GO:0006119). Septic mice at 30 h decreased the transcriptional output of OXPHOS components, as evidenced by a negative GSEA enrichment score (Fig. 1b). Given the connection between oxidative phosphorylation and the tricarboxylic acid (TCA) cycle, we next asked if polymicrobial infection would elicit similar alterations in this pathway. In accordance with the changes observed in the oxidative phosphorylation pathway, we found a negative enrichment score for TCA cycle enzymes (GO:0006099) in the liver of CLP mice (Fig. 1c). Thus, our transcriptome data indicate that sepsis impairs mitochondrial metabolism in the liver.

**Figure 1:**
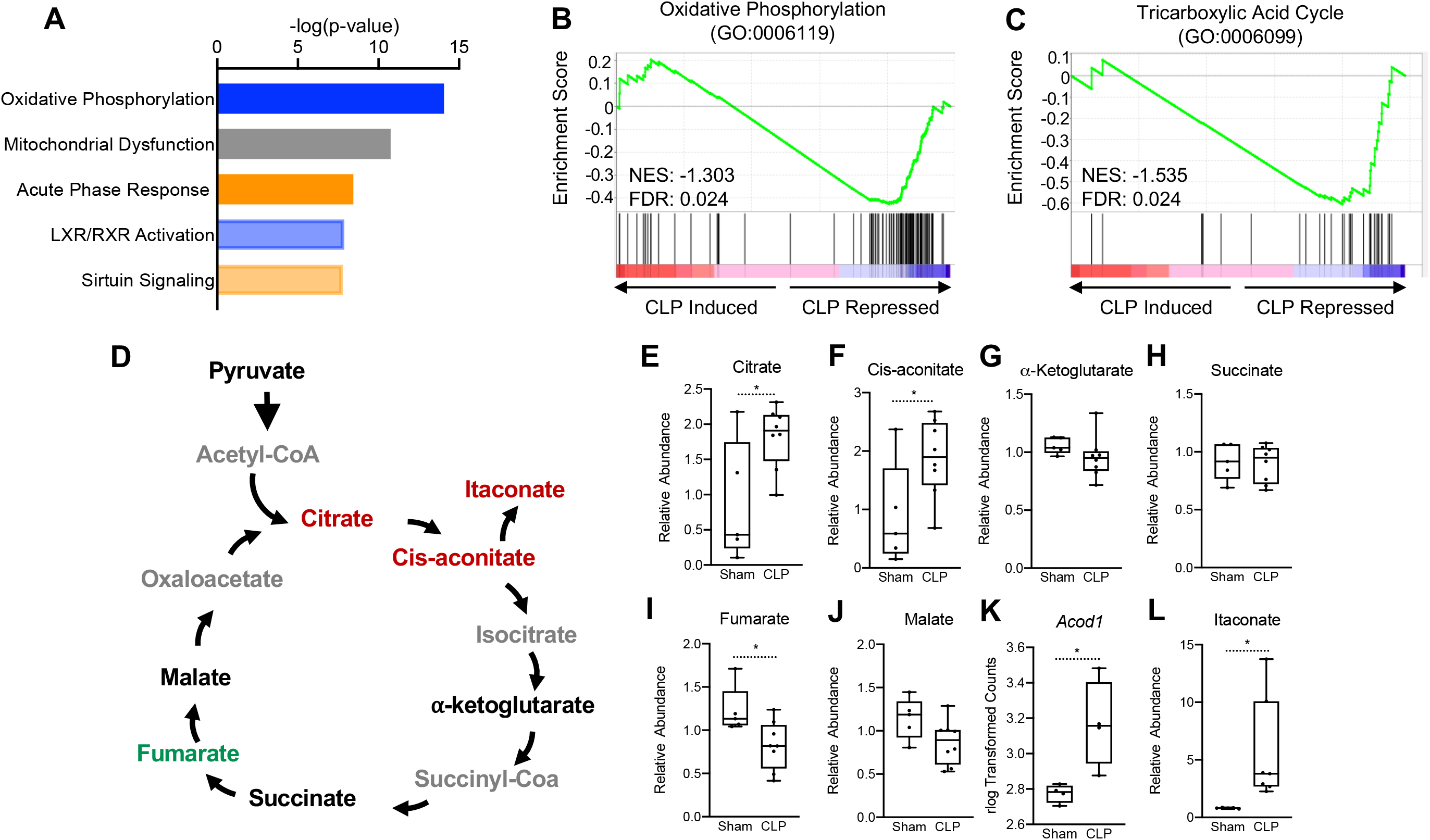
Sepsis Impairs hepatic mitochondrial metabolism. (A) Top 5 canonical pathways subject to transcriptional alterations in the liver identified by Ingenuity Pathway Analysis of RNA-seq of sham versus CLP mice (n = 4 mice per group). Blue represents a negative z-score and orange represents a positive z-score. Shading indicates intensity of pathway activation/inhibition. (B) GSEA of the oxidative phosphorylation pathway (GO:0006119) indicating a negative normalized enrichment score (NES = −0.853). (C) GSEA of the TCA cycle pathway (GO:0006099) indicating a negative enrichment score (NES = −1.1956). (D) Schematic representation of hepatic TCA cycle metabolites altered during chronic sepsis. Red denotes a metabolite increased in response to sepsis; green indicates a metabolite decreased in response to sepsis; black indicates a metabolite unchanged in response to sepsis; grey indicates a metabolite not measured in our metabolomic screening. (E-J) Relative metabolite levels measured by Ultrahigh Performance Liquid Chromatography-Tandem Mass Spectroscopy (UPLC-MS/MS) from livers of sham and CLP mice 30 hours post-surgery (n = 5 sham; 8 CLP). (K) rlog transformed counts from RNA-seq of sham and CLP mice 30 hours post-surgery. (L) Relative itaconate levels in livers of sham and CLP mice measured by UPLC-MS/MS (n = 5 sham; 8 CLP). * denotes p<0.05.

Next, we wanted to assess if the transcriptional changes elicited by sepsis would manifest in altered hepatic TCA cycle metabolism. Therefore, we performed global unbiased metabolomic screening in isolated hepatocytes from control and septic mice by Ultrahigh Performance Liquid Chromotography-Tandem Mass Spectroscopy (UPLC-MS/MS). In line with altered transcriptional regulation of the TCA cycle, we found that sepsis altered the relative abundance of multiple metabolites involved in the TCA cycle (Fig. 1d). In particular, significant elevation of citrate and cis-aconitate was observed in septic hepatocytes at 30h (Fig. 1e&f). Unlike macrophages, which shift their ratio of succinate and a-ketoglutarate to favor succinate accumulation over a-ketoglutarate^29–31^, we found no changes in the levels of these metabolites in hepatocytes (Fig. 1G&H). Furthermore, unlike macrophages^32^, levels of fumarate decreased, but malate levels were unchanged in hepatocytes from septic animals (Fig. 1I&J). Monocytes, macrophages and dendritic cells reprogram the TCA cycle during acute inflammation to a catabolic tolerance phenotype by shunting cis-aconitate to itaconate through the enzymatic action of aconitase decarboxylase (ACOD1; also known as immune responsive gene 1 [IRG1])^32–35^. Accordingly, we also found that sepsis significantly induces *ACOD1* and elevates itaconate in isolate hepatocyte preparations (Fig. 1K&L).

### Sepsis Impairs Hepatic Redox Balance and Promotes Oxidative Stress

Perturbations in oxidative phosphorylation have been shown to promote the production of reactive oxygen species (ROS)^36^. Furthermore, itaconate has been shown to induce genes involved in oxidative stress regulation through activation of the master NRF2 KEAP1 antioxidative pathway in macrophages^34^. Therefore, we examined if sepsis invokes a gene expression profile involved in regulation of ROS. GSEA of the ROS metabolic process pathway (GO:0072593) revealed a positive enrichment, highlighting the transcriptional induction of genes involved in the regulation of oxidative stress in livers of septic mice (Fig. 2a). Given the induction of ROS metabolic genes, we wanted to determine if sepsis alters key metabolites involved in redox balance, specifically the cysteine-glutathione transsulfuration redox regulatory cycle. Sepsis did not alter intracellular homocysteine levels in isolated hepatocytes (Supplemental Fig. 1), however, cystathionine decreased significantly (Fig. 2b). Furthermore, we found a trend for decreased cysteine levels in the liver (*p*=0.0534) (Fig. 2c) and significantly decreased glycine (Fig. 2d). Most dramatically, septic mice depleted hepatocyte glutathione (Fig. 2e) and led to decreases in oxidized glutathione (GSSG) (Fig. 2f). Ophthalamate has recently been shown to be a biomarker of oxidative stress and signals consumption of hepatic GSH^37^. Functionally, we observe a significant accumulation of hepatic ophthalamate levels in response to sepsis, signifying a state of oxidative injury (Fig. 2g).

**Figure 2:**
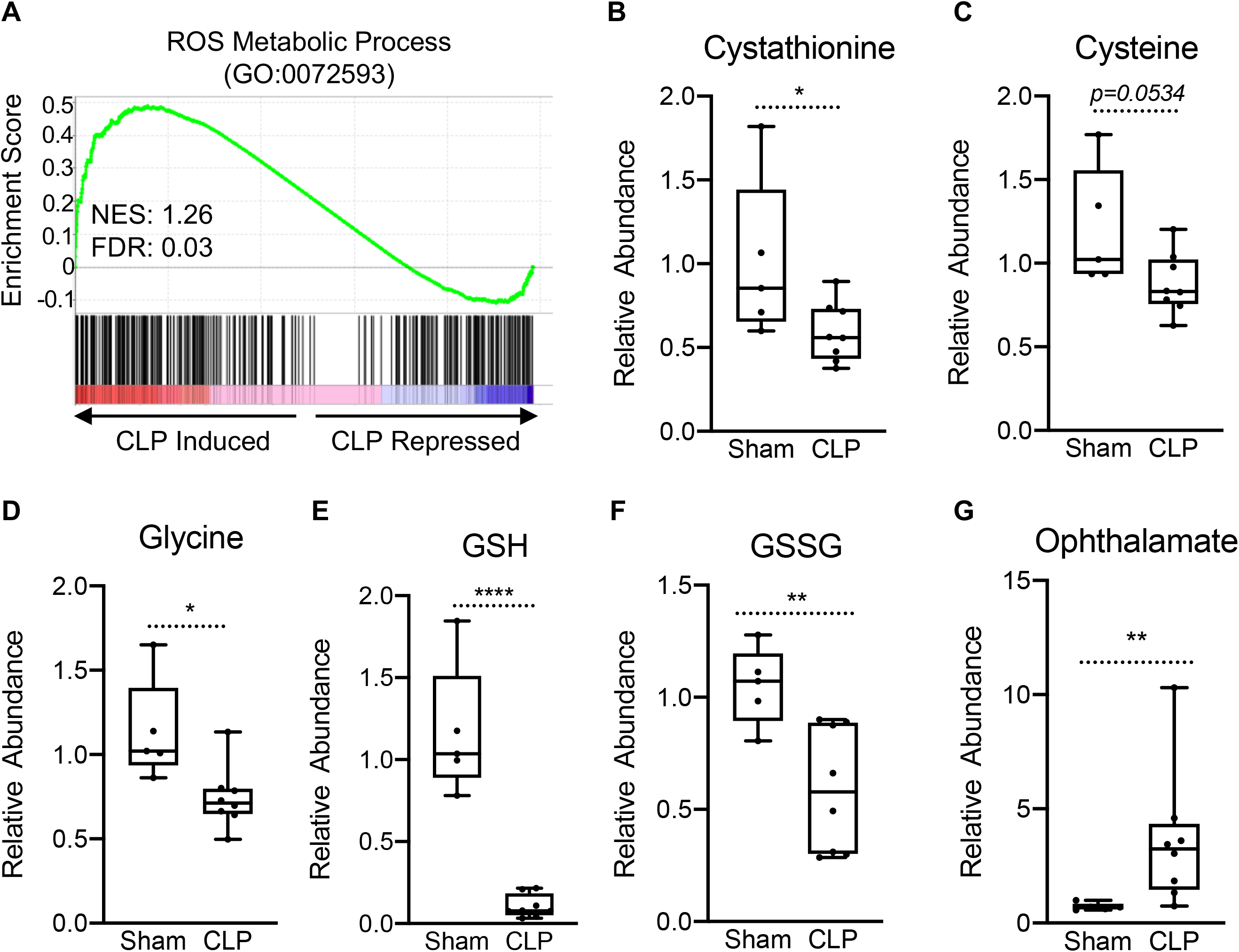
Impaired hepatic redox balance in septic mice. (A) GSEA of the ROS metabolic pathway (GO:0072593) from RNA-seq from livers comparing sham to CLP showing a positive enrichment score (NES = 1.117) (n = 4 mice per group). (B-F) Relative metabolite levels of metabolites involved in redox balance measured by UPLC-MS/MS (n = 5 sham; 8 CLP). (G) Relative levels of ophthalmate involved in redox balance measured by UPLC-MS/MS. * denotes p<0.05, ** p<0.01, **** p<0.0001.

### Sepsis promotes hepatic steatosis

Thus far, our data has revealed that sepsis induces significant impairments in hepatocyte OXPHOS, alterations in the TCA cycle, and induction of oxidative stress, findings that are also metabolic hallmarks of fatty liver disease^38^. Because perturbations in global lipid profiles occur in septic patients^15^, we evaluated the effects of CLP on the hepatic lipid metabolism pathway. In support of our hypothesis that sepsis alters hepatic lipid metabolism, transcriptome data indicate significant repression of lipid metabolism in livers of septic mice (Fig. 3a). Sepsis also decreased transcription output that supports fatty acid metabolic process components in the liver, as evidenced by a negative GSEA enrichment score (Fig. 3b). In particular, we find alterations in genes involved in both fatty acid oxidation as well as fatty acid biosynthesis (Supplemental Fig. 2).

**Figure 3:**
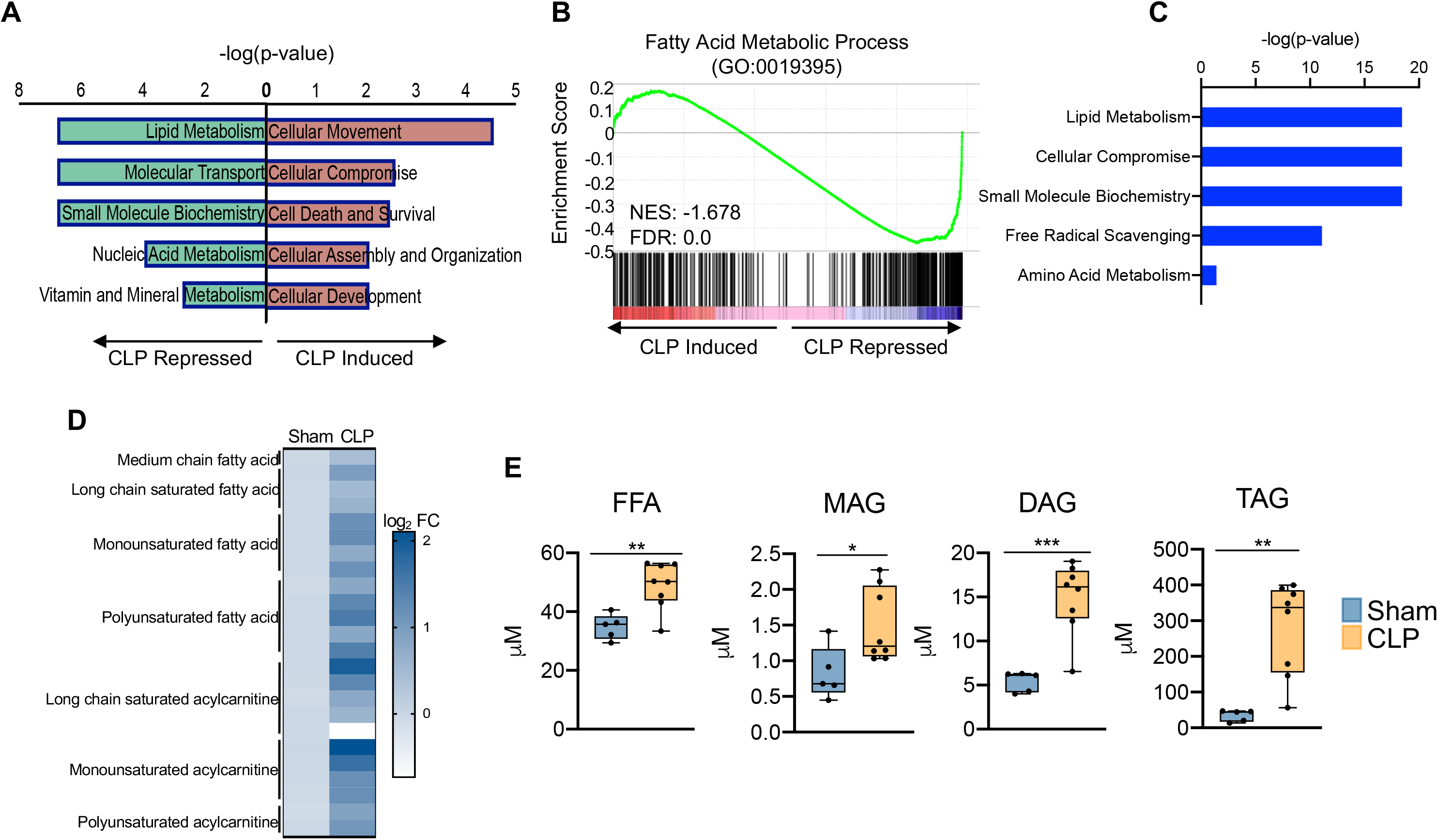
Sepsis promotes hepatic steatosis. (A) Top 5 induced and repressed physiological pathways in the liver of sham versus CLP mice identified by IPA of RNA-seq. (B) GSEA of the fatty acid metabolic pathway (GO:0019395) from RNA-seq from livers comparing sham to CLP showing a negative enrichment score (NES = −1.018) (n = 4 mice per group). (C) IPA of top 5 metabolic pathways significantly altered in the liver in response to sepsis identified by global metabolomic screening. (D) Heatmap representation of log2 fold change of different lipid species in sham and CLP mice measured by UPLC-MS/MS. (E) Quantification of lipids by UPLC-MS/MS in hepatocytes isolated from sham and CLP mice 30 hours post-surgery (n = 5 sham; n = 8 CLP). * denotes p<0.05, ** p<0.01, *** p<0.001.

Since transcriptional regulation of hepatic lipid metabolism is significantly altered during sepsis, we further investigated the functional manifestation of these gene expression profile. Consistent with our transcriptional profiling studies, our unbiased metabolomic screening in hepatocytes also identified lipid metabolism as the top metabolic pathway altered in sepsis (Fig. 3c). We observe increases in virtually all fatty acids and acylcarnitine derivatives assayed (Fig. 3d). The dysregulation in hepatic lipid metabolism in response to sepsis ultimately culminates in the development of steatosis, as evidenced by increased free fatty acids, mono-, di-and triglycerides (Fig. 3e).

### Sepsis remodels the hepatic lipidome

Given the severe dysregulation in hepatic lipid metabolism triggered by sepsis, we next sought to characterize the global consequences on hepatic lipid profiles. This was achieved by targeted lipidomic screening. Parallel to our unbiased metabolomic screening, we find sepsis leads to profound accumulation of almost all lipid species surveyed (Fig. 4a). We next evaluated the hepatic lipid composition to discern which lipid species are must vulnerable to sepsis-induced alterations. We find very little alterations to the ceramide pool in septic hepatocytes (Fig. 4b) with only modest changes in dihydroceramide (Supplemental Fig 3.). However, the hepatic fatty acid fractions displayed significant remodelling in response to sepsis (Fig. 4b). Specifically, we find increases in free fatty acid species such as 16:0, 18:1 and 18:2 in septic hepatocytes (Supplemental Fig. 4a). Morever, hepatic phospholipid composition showed robust alterations following CLP (Fig. 4b). Of the phospholipids (PLs) most significantly affected, we find accumulation of phosphotidylcholines (PCs), phosphotidylethanolamine (PE) and sphingomyelin (SM) (Fig. 4c). Of the hepatic PC pool we find the most abundant species (16:0/18:2 & 16:0/18:1) to show increased accumulion in response to sepsis (Supplemental Fig. 4b). We also observed significant increases in the most abundant PE species (18:0/20:4, 16:0/22:6 & 18:0/22:6) and SM species (22:0, 16:0 & 20:0) to show significant accumulation in septic hepatocytes (Supplemental Fig. 4c&d). We did observe a decrease in one phospholipid species, lysophosphatidylcholine, however, the hepatic concentration of this phospholipid species is signifcantly lower than other PL species assayed (Fig. 4c). Finally, we detected no differences in other PL species such as phosphotidylinositol (PI) and lysophosphatidylethanolamine (LPE) and no differences in hepatic cholesterol ester levels (Supplemental Fig. 5). Our data collectively detail the impact of sepsis on hepatic lipid composition and robustly show a global rise in nearly all lipid species.

**Figure 4:**
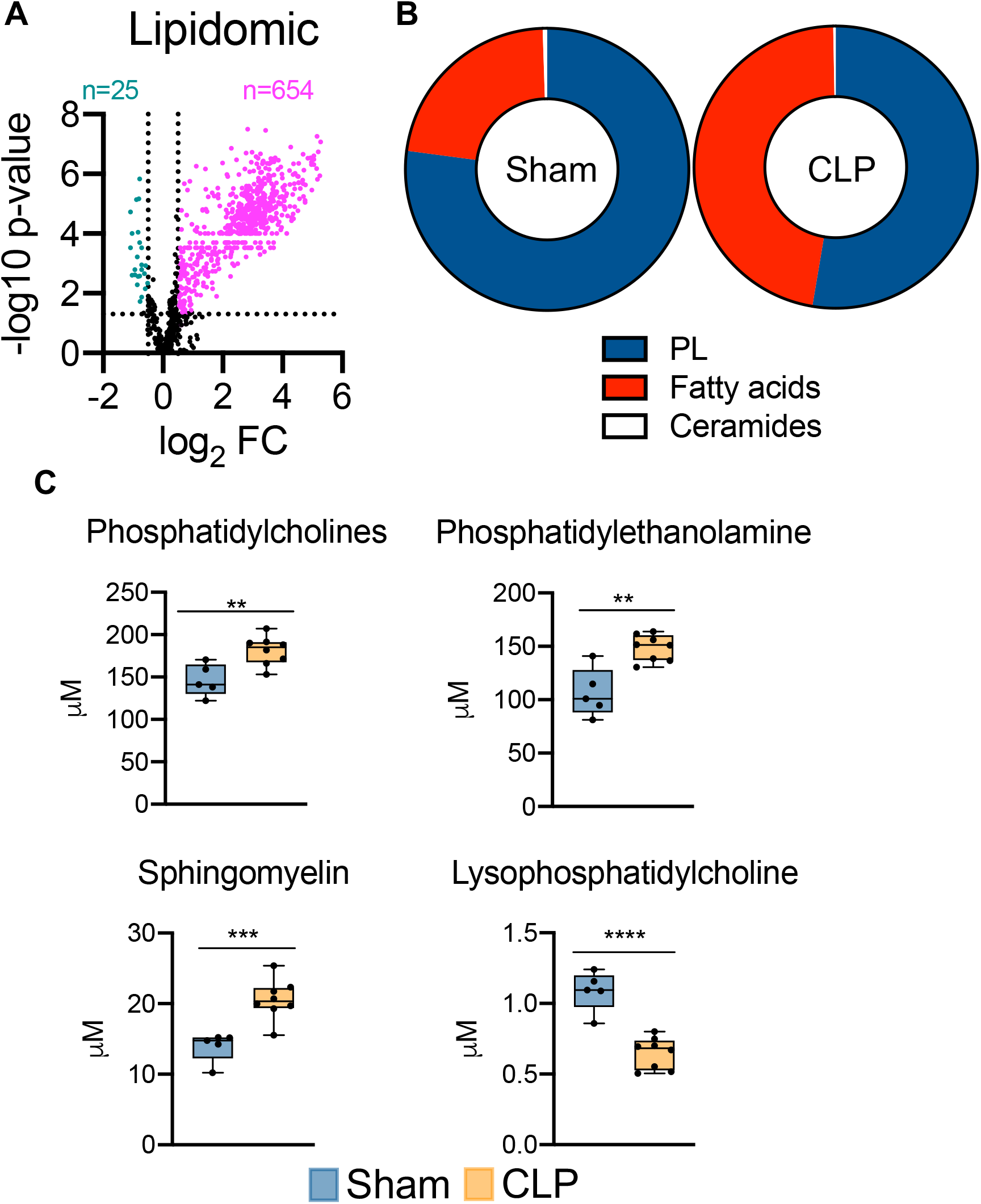
Sepsis reprograms the hepatic lipidome. (A) Volcano plot of significantly altered lipids from sham and CLP hepatocytes measured by UPLC-MS/MS (n = 5 sham; n = 8 CLP). (B) Pie chart of lipid compositions from sham and CLP mice. (C) Quantification of lipids by UPLC-MS/MS in hepatocytes isolated from sham and CLP mice 30 hours post-surgery (n = 5 sham; n = 8 CLP). ** denotes p<0.01, *** p<0.001, **** p<0.0001.

### PDK inhibition attenuates sepsis-induced transcriptional and metabolic changes in the liver

We have previously reported that PDK inhibition by the pyruvate analog and pan-PDK inhibitor dichloroacetate (DCA) promotes PDC-dependent immunometabolic adaptations to sepsis and increases survival^39^. Given the establishment of hepatocyte mitochondrial dysfunction and steatosis during sepsis, we asked whether pharmacological targeting of the hepatic PDK/PDC axis would mitigate the disruption of key metabolic and bioenergetic processes induced by sepsis. To test this postulate, we administered DCA to septic mice 24 h after CLP onset and assessed the global transcription profile of liver 6 h later via RNA-seq. DCA treatment of septic mice reversed the majority of sepsis-regulated gene networks (Fig. 5a). Most notably, pathways involved in redox homeostasis, lipid metabolism and mitochondrial dysfunction are returned to control levels with the administration of DCA in septic mice (Fig. 5b).

**Figure 5:**
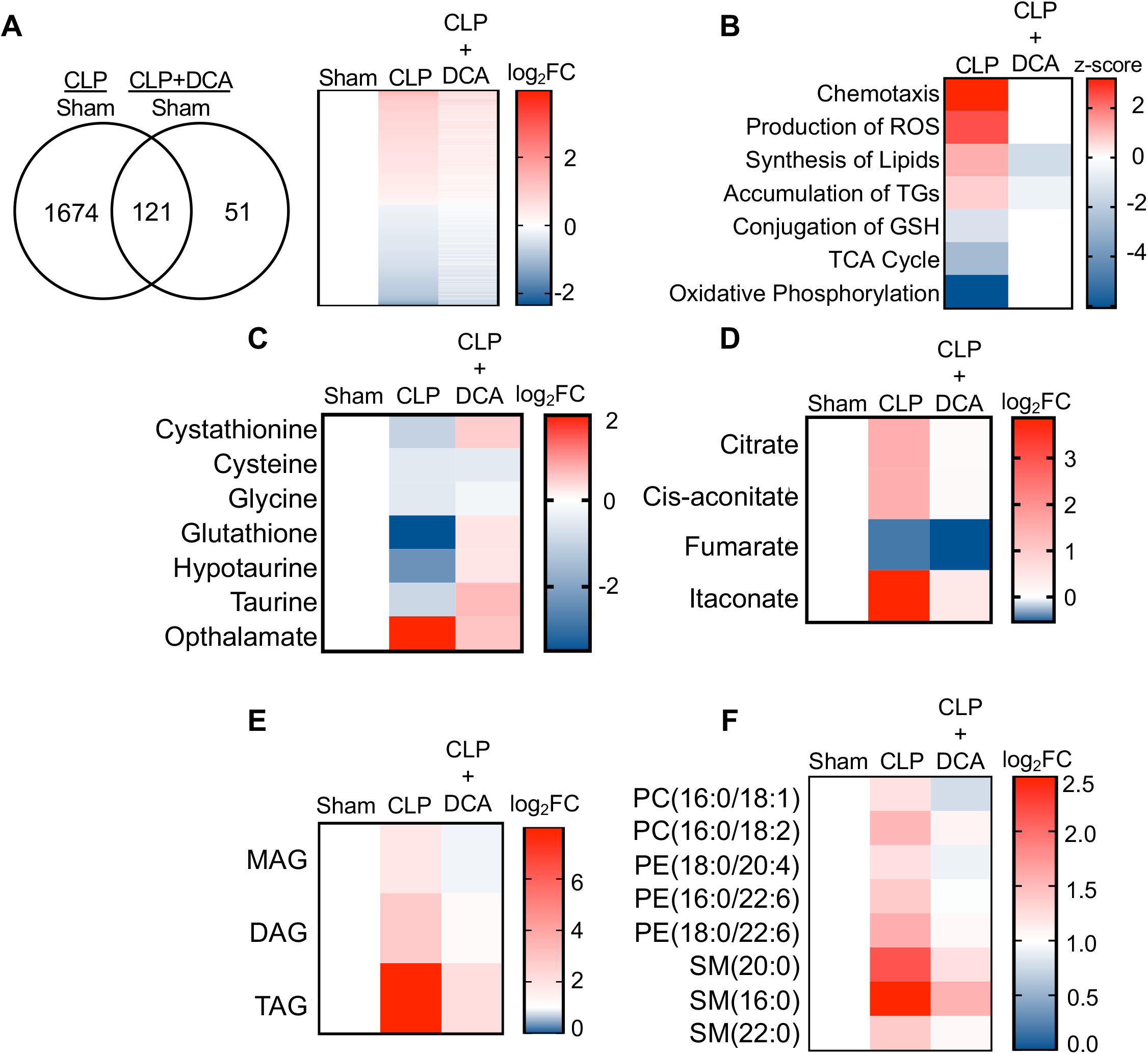
PDK inhibition restores hepatic metabolism in septic mice. (A) Venn diagram of differentially expressed genes (DEGs) assessed by RNA-seq in CLP versus sham compared to CLP+DCA versus sham 30 hours post-surgery (left). Heatmap of average log2 fold change of DEGs in sham, CLP and CLP+DCA (right) (n = 4 mice per group). (B) Heatmap depiction of z-scores of top canonical pathways identified by IPA in CLP versus sham and CLP+DCA versus sham. (C) Heatmap depiction of average log2 fold change in metabolite levels involved in redox balance in sham, CLP and CLP+DCA 24 hours post-surgery measured by UPLC-MS/MS (n = 5 sham; n = 8 CLP and CLP+DCA). (D) Heatmap depiction of average log2 fold change in metabolite levels involved in TCA cycle in sham, CLP and CLP+DCA 30 hours post-surgery measured by UPLC-MS/MS (n = 5 sham; n = 8 CLP and CLP+DCA). (E) Heatmap depiction of average log2 fold change in MAG, DAG and TAG levels in sham, CLP and CLP+DCA 30 hours post-surgery measured by UPLC-MS/MS (n = 5 sham; n = 8 CLP and CLP+DCA). (F) Heatmap depiction of average log2 fold change in phospholipids in sham, CLP and CLP+DCA 30 hours post-surgery measured by UPLC-MS/MS (n = 5 sham; n = 8 CLP and CLP+DCA.

Given our previous results indicating the transcriptional changes associated with sepsis in the liver underlie functional alterations in metabolism, we wanted to determine if the DCA reversal of transcriptional changes mounts to differential metabolomes. Given our RNA-seq data showing a positive enrichment score for production of ROS along with negative enrichment for glutathione conjugation, we started off with evaluating the redox pathway metabolites. Strikingly, the depletion of redox components cystathionine, cysteine, glycine, hypotaurine, taurine and glutathione elicited by sepsis was completely reversed by administration of DCA (Fig. 5c). More importantly, we observed a significant decrease in hepatic ophthalmate levels, signifying a functional decrease in oxidative stress (Fig. 5c). The restoration in redox metabolites in septic mice following DCA administration prompted us to investigate the mitochondrial consequences of this pharmacological intervention. With the exception of fumarate, DCA completely restores TCA metabolite levels to that of sham levels (Fig. 5d). Finally, we wished to define the effects of DCA treatment on the hepatic lipidome. Lipidomic analysis shows a protective effect of DCA in ameliorating sepsis-induced steatosis. In sum, these data reveal pharmacological inhibition of the PDK pathway restores transcriptional and metabolic changes associated with sepsis in the liver.

## Discussion

Considerable attention has been focused recently on the immunometabolic consequences of inmflammation in immune cells^40^. However, significant gaps exist in our understanding of the metabolic adaptations induced by systeminc inflammation from sepsis in vital organs. Addressing this limitation is critical because sepsis survival depends on restoring both organ and immune cell homeostasis following dysregulated and disseminated inflammation^41^. In the present study, we reveal severe disruption of several key hepatic metabolic pathways in septic mice, including TCA cycle activity, OXPHOS, redox balance and lipid metabolism. Importantly, DCA administration restored TCA cycle flux, restored rerdox balance and ameliorated sepsis-induced steatosis. Our study highlights the clinical potential of targeting the PDC/PDK axis in restoring liver function/metabolism in the context of sepsis.

Remodelling of TCA cycle flux is well appreciated to support immune cell effector function in the face of inflammatory challenges^3,31,42,43^. For example, succinate primes inflammation through succinate dehydrogenase (SDH)-mediated ROS generation and ATP synthesis^31,42^. SDH-derived ROS enhances IL-1ß production by activating HIF-1a and activating the NLRP3 inflammasome^20^. Inhibition of SDH activity increases IL-10 production and skews inflamed immune cells towards an anti-inflammatory response^31^. Fumarate, on the other hand, possesses anti-inflammatory properties as it inhibits pro-inflammatory cytokine production either by activating the NRF2 pathway or via inhibiting pathways like NF-κB and MAPK that lie downstream of TLR signaling in immune cells^44,45^. Our findings extend the concept that the TCA cycle is subject to reprogramming in response to sepsis in non-immune cells such as hepatocytes. Furthermore, we found similarities as well as differences in terms of TCA cycle remodelling in hepatocytes compared to immune cell effects, clearly indicating this pathway is regulated in a cell-type and context-dependent manner during sepsis.

In response to LPS stimulation, macrophages significantly upregulate IRG1 and accumulate itaconate from the decarboxylation of cis-aconitate^32,34^. One of the primary functions itaconate serves in immune cells is to limit IL-1 β production through direct inhibition of SDH and activation of the NRF2/Keap1 and IkBζ-ATF3 pathways^32,34,46^. In line with what occurs in macrophages, we observed an increase in IRG1 and itaconate in septic hepatocytes at 30h. While we did not fully characterize the hepatic function of itaconate, a recent paper demonstrates its anti-inflammatory effects during Ischemiareperfusion (IR) injury in the liver^47^. The deletion of IRG1 heightens inflammation and liver damage and renders hepatocytes susceptible to oxidative injury after I/R injury. Further, itaconate administration reduces liver damage and inflammation associated with I/R in IRG1 KO mice, also emphasizing its anti-inflammatory and hepatoprotective effects^47^. Given the immunomodulatory effects of itaconate and its accumulation in the liver in our study, we hypothesize that itaconate directs hepatocyte shifts in both TCA cycle metabolism and mitochondrial energetics. DCA reduces itaconate levels, which we also find in a human monocyte model of sepsis^48^.

Similar to immune cells, liver cells increase ROS production during sepsis^49,50^. We offer a potential mechanism whereby sepsis leads to significant depletion of redox metabolites within the glutathione pathway ultimately culminating in loss of cellular glutathione levels resulting in ophthalamate accumulation in hepatocytes.

Sepsis profoundly alters lipid metabolism, resulting in significant increases in plasma fatty acid and glycerol concentrations, changes reported to predict prognosis in septic patients^51^. Furthermore, patients who died from sepsis had evidence of hepatic steatosis affecting 5%-80% of liver parenchyma^52^. Human data supports a robust hepatic steatosis phenotype subsequent to sepsis. One vital lipid metabolic pathway which can contribute to steatosis is *de novo* lipogenesis. While we did not directly measure hepatic lipogenesis, our metabolomics screening did show an increase in hepatic citrate and free fatty acid levels suggesting an increased lipogenic drive during sepsis. Functionally, immune cells exploit the lipogenic pathway for the generation of malonyl-CoA and subsequent malonylation of glycolytic enzymes, which ultimately contributes to the metabolic adaptation of macrophages to inflammatory cues^53^. Future studies dissecting the role of lipogenesis in altering hepatocyte function during systemic infection are warranted.

In addition to accumulation of free fatty acids and triglycerides in the liver, our lipidomic analysis revealed significant increases in phospholipids. In the present study, we find hepatic levels of the phospholipids phosphatidylcholine, phosphatidylethanolamine, and sphingomyelin to be heightened during sepsis, indicating abnormal metabolism of phospholipids. PC and PE are the most abundant phospholipids that make up cellular membranes and an abnormally high ratio of PC and PE has been reported to influence energy metabolsm and linked to fatty liver disease and impairement in liver regeneration after injury^54–56^. Altered PC: PE ratio has also been shown to influence the dynamics and regulation of lipid droplets contributing towards steatosis, a phenotype we also report in our study^54,55,57^. The dysregulation of hepatic phospholipid metabolism may be a contributing mechanism underlying steatosis, which is a topic of future investigation. Additionally, phospholipids serve as precursor molecules to bioactive lipids, which are involved in numerous signal transduction cascades^58^. Investigating whether signaling properties of bioactive lipids in the liver during sepsis is pathogenic or contributes to sepsis resolution is also needed in the future.

PDC is a master metabolic regulator controlling the conversion of pyruvate to acetyl-CoA in the mitochondria^59^. A part of the metabolic reprogramming that occurs in immune cells in response to inflammatory signaling is inactivation of PDC^22,59^. PDK is a negative regulator of the PDC, as it phosphorylates PDC and inhibits the conversion of pyruvate to acetyl-CoA^59^. During sepsis, the expression and activity of PDK1 in immune cells is heightened, contributing to the dysfunction of mitochondria metabolism. One mechanism driving this observed increase is through pro-inflammatory mediators such as LPS and interferon gamma (IFN-γ)^60^. Another possible mechanism contributing to PDK activation in the context of sepsis is through glucocorticoid signaling^61^. In fact, starvation is well established to activate the PDK pathway^62–66^. Based on our findings of elevated glucocorticoid levels in both the circulation and the liver, we hypothesize the stress hormone pathway as a contributor to PDK activation during sepsis.

We reported in a sepsis monocyte model that DCA reduced TCA cycle catabolic effects concurrent with increasing amino acid anaplerotic catabolism of branched-chain amino acids, leading to increased TCA-driven anabolic energetics^22^. In the present study, we find restoration of hepatic TCA metabolites, decreased triglyceride accumulation, lessening of lipid synthesis and oxidative stress rebalances in septic mice after DCA administration. Overall, we demonstrate that hepatic transcriptional and metabolic dysfunction improves after targeting the PDC/PDK axis with DCA.

Further work elucidating the mechanistic pathways involved in the dysfunction of lipid metabolism and the impact of TCA metabolite alterations is warranted. In particular, further investigation specifically into the PDK pathway to determine which isoform underlies the hepatic manifestions of sepsis. This is warranted given multiple isoforms of PDK expressed in the liver^67^ and DCA inhibits multiple isoforms of PDK^68,69^. The results of this study fill a gap in understanding how sepsis at the molecular level compromises liver function. It also further informs the novel therapeutic targeting potential of the PDC/PDK homeostat.

## Material and Methods

### Animal Experiments

Male C57BL/6J mice aged 8-10 weeks were purchased from The Jackson Laboratory (Bar Harbor, ME). All animals were subject to a 12:12 hour dark/light cycle with *ad libitum* access to standard rodent chow and water. Cecal ligation puncture model was used to induce sepsis as previously described^39,70^. Briefly, cecum was ligated and punctured two times with a 22-gauge needle. Contents were then returned and incision was closed in two layers (peritoneum and skin). Sham operation where abdominal incision was made but cecum not ligated or punctured was used as a control. Subcutaneous fluids (1ml normal saline) were given to each animal. Mice were euthanized 30 hours post-surgery for tissue collection. Dichloroacetate (DCA) (Sigma; MO, USA) was administered (25mg/kg) intraperitoneally at various time points post-surgery: for metabolomic screening and RNA-seq DCA was administered 24 hours post-surgery and tissues collected 6 hours post DCA administration (30 hours post-surgery), for metabolic cages DCA was administered immediately post-surgery.

### Hepatocyte Isolation

Hepatocytes were isolated via portal vein perfusion and collagenase digestion as previously described^71^. Following perfusion, liver cells were liberated by gentle dissocation in DMEM (ThermoFisher; CA, USA). Cells were then filtered through nylon mesh to remove cellular debris and connective tissue and resulting cells pelleted by centrifugation at 50g for 1 minute. After three washes with DMEM cells were counted and viability assessed via Trypan Blue exclusion.

### RNA-sequencing

RNA was isolated from whole liver tissues using Trizol and the RNeasy RNA isolation kit (Qiagen; MD, USA) according to manufacturer’s protocol. One microgram of high quality RNA (RIN>8) was used as a template for library generation using the Illumina TruSeq RNA Sample Prep Kit v2 (Illumina; CA, USA) according to the manufacturer’s protocol. Generated libraries were then poly(A) enriched for mRNA prior to sequencing. Indexed samples were sequenced at 100bp-paired-end protocol with the NovaSeq 6000 (Illumina), generating approximately 20-30 million reads per sample. Sequenced reads were aligned to the University of California Santa Cruz (UCSC) mm10 reference genome using STAR v2.5 as previously described^72^. The mapped read counts were quantified by Subread feature Counts v1.5.0-p1^73^. Differentially expressed genes (DEGs) were determined by DESeq2 v1.14.1^73^ using a false discory rate of 0.05. Ingenuity Pathway Analysis (Qiagen) and Gene Set Enrichment Analysis v4.0.3 (GSEA) were further used as previously described^74^.

### Ultrahigh Performance Liquid Chromatography-Tandem Mass Spectroscopy

Hepatocytes were isolated described above for metabolomic screening via Ultrahigh Performance Liquid Chromotography-Tandem Mass Spectroscopy (UPLC-MS/MS) (Metabolon; NC, USA). Briefly, 150-200 microliter cell pellets per animal were used as starting material. Samples were prepared using the automated MicroLab STAR system (Hamilton; NV, USA). Proteins were precipitated using methanol under vigorous shaking for 2 minutes followed by centrifugation. Prior to analysis organic solvents were removed with TurboVap (Zymark; MA, USA) and overnight storage under nitrogen. Dried samples were reconstituted with solvents compatible with the three following analytical methods: 1) reverse phase (RP)/UPLC-MS/MS methods with positive ion mode electrospray ionization (ESI), 2) RP/UPLC-MS/MS with negative ion mode ESI and 3) HILIC/UPLC-MS/MS with negative ion mode ESI. Resulting samples were analyzed with the ACQUITY UPLC (Waters; MA, USA) and a Q-Exactive high resolution/accurate mass spectrometer (ThermoScientific; MA, USA) interfaced with a heated electrospray ionization (HESI-II) source and Orbitrap mass analyzer operated at 35,000 mass resolution. An acidic positive ion condition was used on an aliquot optimized to detect more hydrophilic compounds. Another acidic positive ion condition was ran but chromotographically optimized for hydrophobic compounds. Basic negative ion optimized conditions was also used on a separate C18 column. Resulting raw data was extracted and peaks identified using Metabolon’s hardware and software. Compounds were identified by comnparing to known library entries of purified standards or recurrent unknown entities. A lirary of authenticated standards containing retention time/index (RI), mass to charge ratio *(m/z)* and chromotographic data (MS/MS spectral data) on all library compounds. Three criteria are used to identify chemicals: 1) RI within a narrow window of proposed identification, 2) accurate mass match to the library ± 10 ppm and 3) MS/MS forward and reverse scores between the experimental data and authentic standards. Peaks were quantified using area under the curve (AUC). For complex lipid panel lipids were extracted with methanol:dichloromethane in the presence of internal standards. Extracts were concentrated under nitrogen and reconstituted in 250ml of 10mM ammonium acetate dichloromethane:methanol (50:50). Mass spec analysis was performed in a Shimazdu LC with nano PEEK tubing and the Sciex SlexIon-5500 QTRAP (Sciex; MA, USA). Both negative and positive mode electrospray was used. Individual lipid species were quantified by taking the peak area ratios of target compounds and their assigned internal standards then multiplying by the concentration of added internal standards. Lipid class concentrations were calculated from the sum of all molecular species within a class, and fatty acid compositions were determined by calculating the proportion of each class comprised by individual fatty acids.

## Acknowledgements

The authors of this manuscript would like to thank the National Institute of Environmental Health Sciences Epigenomics and DNA Sequencing Core for their help with RNA-sequencing and Dr. Sara Grimm for her analysis of transcriptome data. This work was supported by NIH Intramural Program 1ZIAES10328601 (J.M.), NIH R01 HL132035 (X.Z.), NIH R01 HL119962 (J.S.P.), NIH K01 DK117069 (C.K), NIH K01 AG056663 (E.Q.), NIH R01 AI065791 (C.E.M.), R01 GM102497 (C.E.M.), R35 GM126922 (C.E.M.).

---

**Supplemental figure 1:**
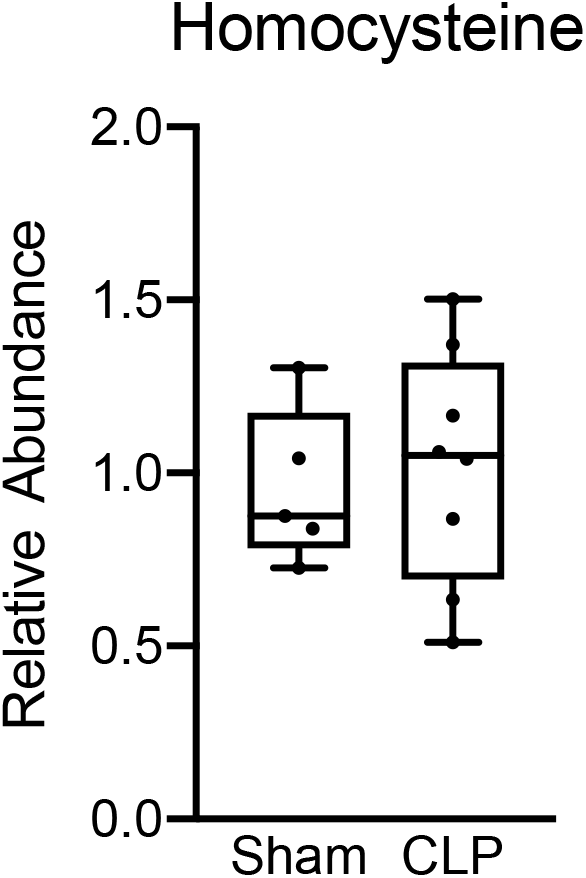
Hepatic homocysteine levels in sham and CLP mice. Ultrahigh Performance Liquid Chromatography-Tandem Mass Spectroscopy measurement of homocysteine from isolated hepatocytes 30 hours post-surgery. n=5 sham, n=8 CLP.

**Supplemental figure 2:**
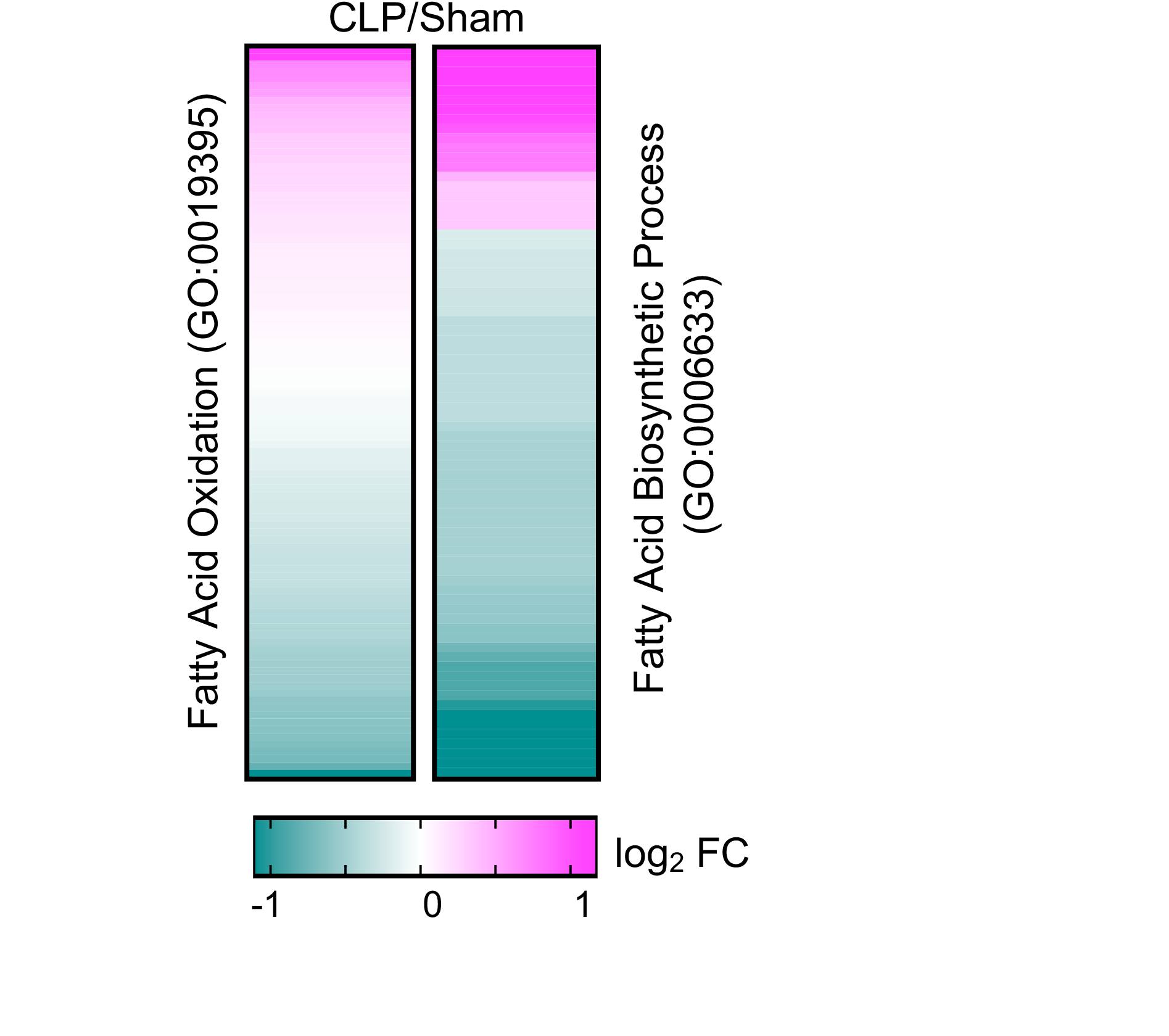
Fatty acid oxidation and biosynthetic pathways in septic livers. Heatmap representation of average log2 fold change of genes involved in fatty acid oxidation (GO:0019395) and fatty acid biosynthetic process (GO:0006633) in CLP versus sham operated mice (n = 5 sham; n = 8 CLP).

**Supplemental figure 3:**
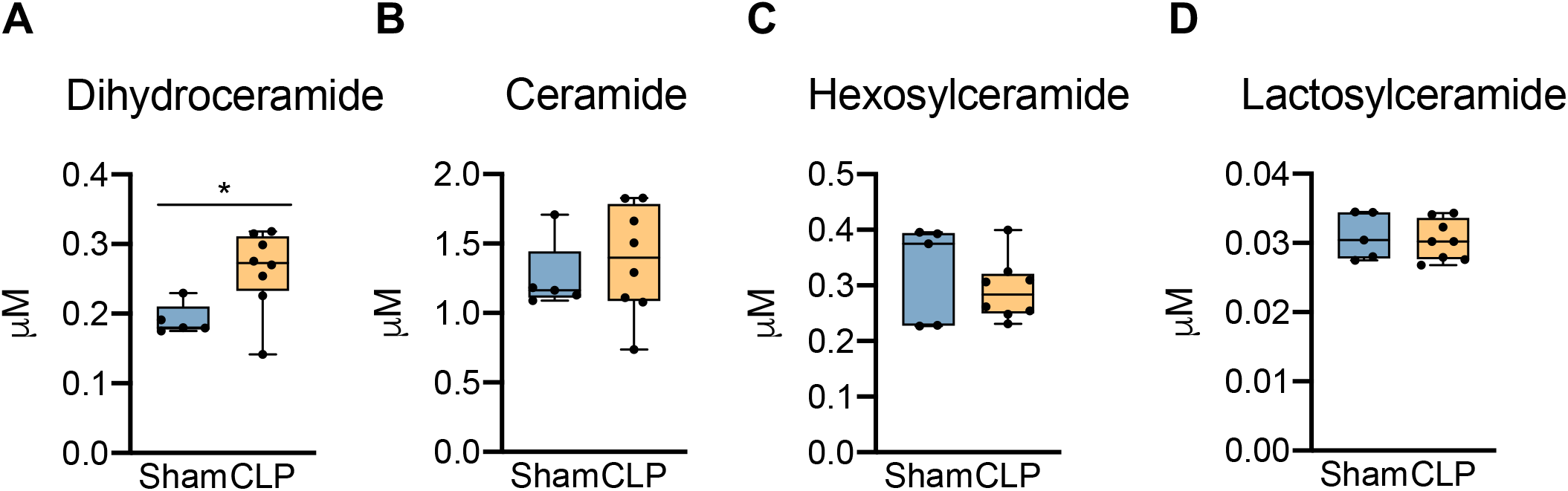
Hepatic ceramide levels in sham and CLP mice. Ultrahigh Performance Liquid Chromatography-Tandem Mass Spectroscopy measurement of **(A)** dihydroceramide, **(B)** ceramide, **(C)** hexosylceramide and **(D)** lactosylceramide from isolated hepatocytes 30 hours post-surgery. n=5 sham, n=8 CLP. * denotes *p*<0.05 determined by Student’s T-test.

**Supplemental figure 4:**
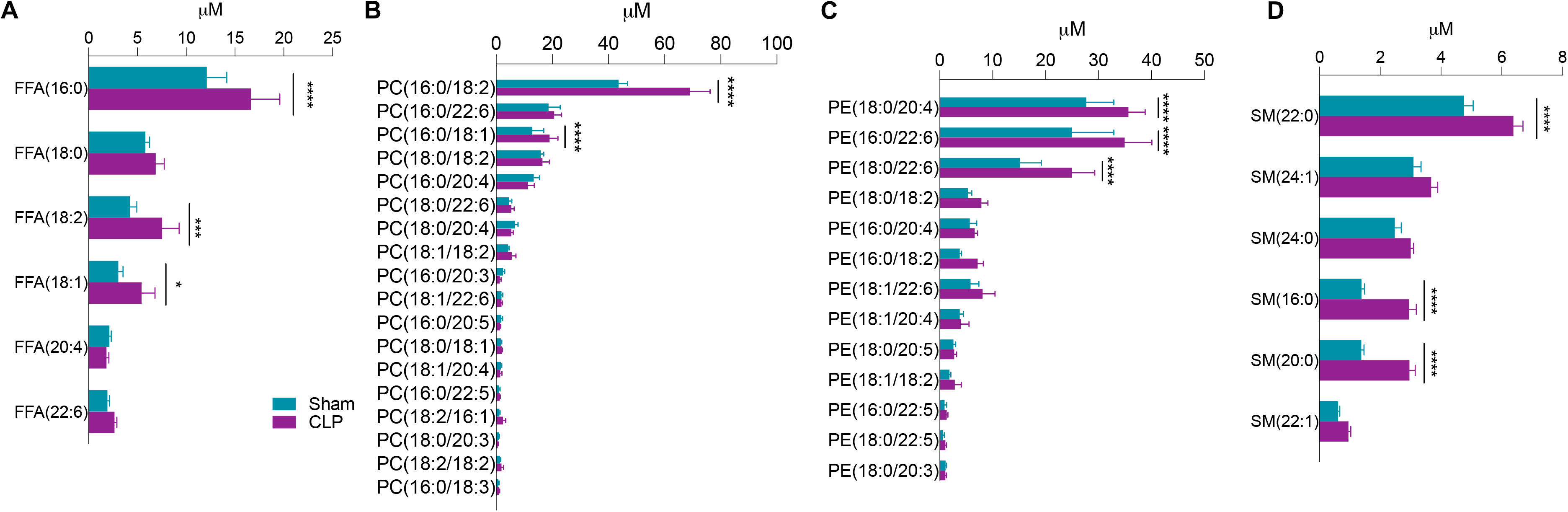
Phosphatidylcholine, phosphatidylethanolamine, and sphingomyelin quantifications. Ultrahigh Performance Liquid Chromatography-Tandem Mass Spectroscopy measurement of **(A)** free fatty acid species, **(B)** phosphatidylcholine species, **(C)** phosphatidylethanolamine species and **(D)** sphingomyelin species from isolated hepatocytes 30 hours post-surgery. n=5 sham, n=8 CLP. **** denotes *p*<0.0001 determined by two-way ANOVA followed by multiple comparisons test.

**Supplemental figure 5:**
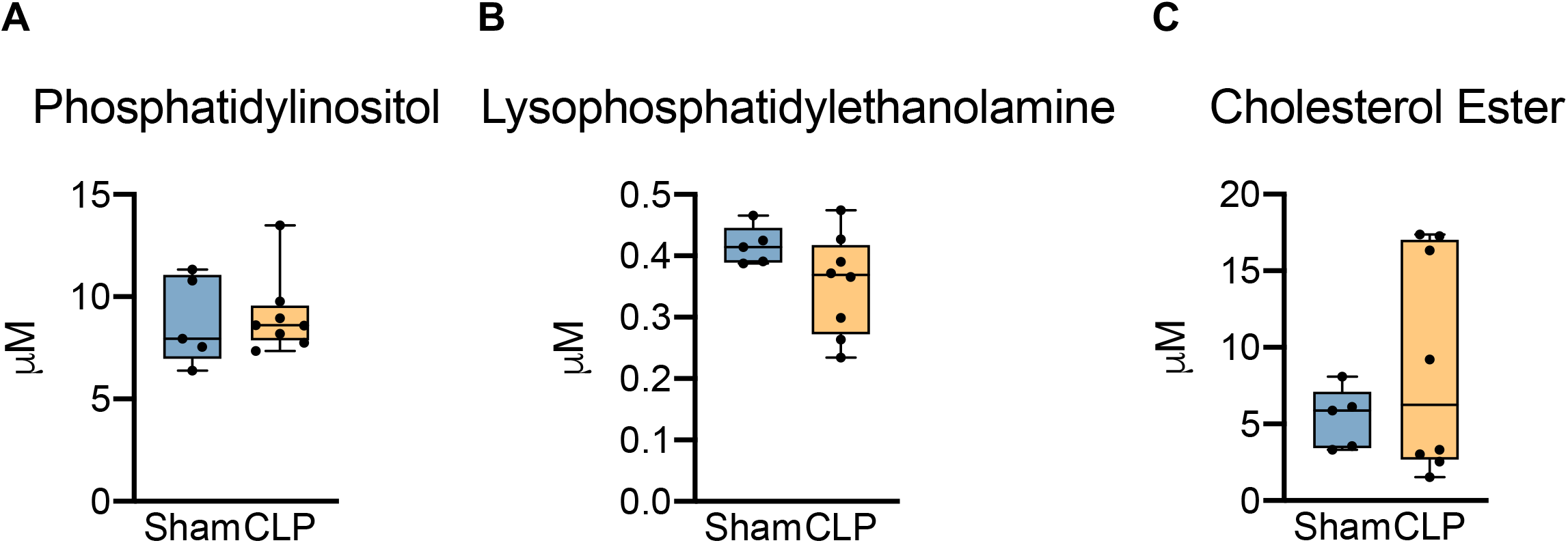
PI, LPE and CE levels in septic livers. Ultrahigh Performance Liquid Chromatography-Tandem Mass Spectroscopy measurement of **(A)** phosphatidylinositol, **(B)** lysophosphatidylethanolamine species and **(C)** cholesterol ester from isolated hepatocytes 30 hours post-surgery. n=5 sham, n=8 CLP. **** denotes *p*<0.0001 determined by two-way ANOVA followed by multiple comparisons test.

